# MolDeBERTa: Foundational Model for Physicochemical and Substructure-Informed Molecular Representation Learning

**DOI:** 10.64898/2026.02.15.706011

**Authors:** Gabriel Bianchin de Oliveira, Fahad Saeed

**Affiliations:** Knight Foundation School of Computing and Information, Sciences, Florida International University, Miami, Florida, USA

**Keywords:** Molecular Foundational Models, Representation Learning, Transformer Encoders, Chemical Language Models, SMILES

## Abstract

Foundational models that learn the “language” of molecules are essential for accelerating material and drug discovery. These self-learning models can be trained on large collections of unlabelled molecules, enabling applications such as property prediction, molecule design, and screening for specific functions. However, existing molecular language models rely on masked language modeling, a generic token-level objective that is agnostic to physico-chemical and substructure molecular properties. Here we introduce *MolDeBERTa*, a chemistry-informed self-supervised molecular encoder built upon the DeBERTaV2 architecture with byte-level Byte-Pair Encoding (BPE) tokenization. MolDeBERTa is pre-trained on up to 123 million SMILES from PubChem using three novel pretraining objectives designed to inject strong inductive biases for molecular properties and substructure similarity directly into the latent space. The model is systematically investigated across three architectural scales, two dataset sizes, and five distinct pretraining objectives, of which three are novel and two are adapted from prior work. When evaluated on 9 MoleculeNet benchmarks, MolDeBERTa achieves the best overall performance on 4 out of 9 tasks and outperforms SMILES-based encoders on 7 out of 9 tasks, with up to a 16% reduction in regression error, and improvements of up to 2.2 ROC-AUC points on classification tasks. The source code, pretrained checkpoints, and datasets are publicly available at https://github.com/pcdslab/MolDeBERTa and https://huggingface.co/collections/SaeedLab/moldeberta.

**CCS Concepts:**

- **Computing methodologies** → **Neural networks**; **Natural language processing**; **Learning paradigms**.

## 1 Introduction

Foundational models have become a central paradigm for molecular representation learning, enabling data-driven discovery across chemistry [5], materials science [25], and drug development [3]. By learning from large collections of molecular sequences such as SMILES, these models have demonstrated strong performance across a wide range of downstream tasks, including molecular property prediction [17], protein-small molecule interaction modeling [12], and binding affinity estimation [8].

Masked Language Modeling (MLM) has been the dominant pretraining strategy for molecular encoders used in methods such as ChemBERTa [6] and MolFormer [23]. These methods have demonstrated that large-scale pretraining improves performance across a wide range of tasks, and can learn meaningful molecular representations [1, 6, 16, 23, 26, 31]. More recent work includes usage of large decoder-based generative language models [5, 15, 30] capable of molecule generation and autoregressive reasoning.

Despite these advances, current molecular representation learning remains limited for discriminative tasks. Decoder-based generative models are optimized for sequence generation and unidirectional prediction, making them computationally expensive and less suitable for tasks that require bidirectional representations [4]. At the same time, encoder-based molecular models have seen limited evolution in pretraining methodology, with most approaches relying on MLM objectives that operate at the token level and do not explicitly encode molecular properties or substructure information. These models are typically built on transformer architectures developed for natural language processing, such as RoBERTa [18] and BERT [7], whose pretraining objectives were designed for syntactic and semantic linguistic patterns rather than chemical ones. Consequently, molecular encoders trained with generic token-level objectives tend to learn representations that are less aligned with chemical properties, limiting their effectiveness in downstream discriminative tasks [1, 22, 27].

Beyond encoder-based language models, the molecular representation learning landscape includes methods that leverage richer molecular modalities. Graph-based approaches, such as those based on message-passing neural networks, operate directly on 2D molecular graphs, and have demonstrated strong performance across property prediction tasks [22, 29]. More recently, 3D-aware models that incorporate geometric information, such as atomic coordinates, and conformational properties, have shown further improvements on tasks where spatial structure is critical [10, 32]. Graph-based methods, while effective, can introduce additional computational overhead through explicit graph construction and specialized architectures. In addition, the 3D-aware models require pre-computed conformers that may be unavailable for the vast majority of compounds in large-scale databases. SMILES-based encoders offer a simpler and more scalable pipeline, making them particularly attractive for high-throughput settings and for practitioners without access to specialized molecular modeling infrastructure.

In this work, we designed and developed *MolDeBERTa*, a family of encoder-based molecular foundational models that leverages the DeBERTaV2 architecture with chemistry-informed pretraining objectives that explicitly guide the representation space toward molecular properties and substructure similarity, *without* relying on human-annotated labels or experimental data. We systematically study the interaction between pretraining objective, model scale, and dataset scale across 30 pretrained variants evaluated on nine MoleculeNet benchmarks. Our results show that chemistry-informed pretraining substantially improves representation quality, outperforming SMILES-based encoders on 7 out of 9 tasks, and achieving the best overall performance on 4 out of 9 tasks across all evaluated paradigms, with improvements up to 2.2 ROC-AUC and 16% RMSE reduction. Our contributions in this paper can be summarized as follows:

- We introduce *MolDeBERTa*, a DeBERTaV2-based encoder foundational model for SMILES-based molecular representation learning. Our contribution lies in the systematic adaptation of a stronger encoder backbone combined with chemistry-informed pretraining objectives.
- We propose physicochemical and substructure-informed pretraining objectives beyond MLM, including multi-label substructure classification, and descriptor-informed contrastive learning, explicitly aligning the latent space with chemical properties without requiring experimental labels or 3D geometry.
- We conduct a large-scale study across 30 pretrained models, analyzing the interaction between pretraining objective, model scale, and dataset scale on nine MoleculeNet benchmarks. To the best of our knowledge, this is the most comprehensive ablation study for encoder-based SMILES foundational models.
- We provide scientific validation through atom-level attribution analyses, showing that MolDeBERTa learns task-specific representations aligned with known physicochemical principles and outperforms state-of-the-art models on downstream property prediction tasks.

## 2 Related Work

### 2.1 Encoder-based Foundational Models for SMILES

Recent advances in molecular representation learning have demonstrated that encoder-based foundational models pretrained on SMILES strings can learn transferable representations for a wide range of chemical tasks. Early approaches, such as SMILES-BERT [26], MTL-BERT [31], and Mol-BERT [16], adapted BERT-based masked language modeling to molecular strings, establishing the feasibility of treating SMILES as a chemical language. Subsequent models, including ChemBERTa-1 [6] and its later variants [1], scaled this paradigm by pretraining on larger molecular corpora and incorporating alternative objectives such as multi-task regression over molecular descriptors. MolFormer [23] introduced architectural modifications and large-scale pretraining to further improve molecular performance, demonstrating results across MoleculeNet [28] benchmarks.

While these encoder-based models have demonstrated the effectiveness of SMILES representation learning, they largely rely on transformer architectures developed for natural language processing, such as BERT [7] and RoBERTa [18]. Advances in encoder-based transformer architectures for natural language processing, including the DeBERTa family [9], have shown improved representation learning capabilities, but have not yet been explored in the context of molecular language models. As a result, the potential benefits of modern encoder architectures for molecular representation learning remain largely unexplored.

### 2.2 Pretraining for Molecular Representation

The choice of pretraining objective plays an important role in determining the quality of molecular representations. Masked Language Modeling (MLM) remains the dominant pretraining objective for SMILES-based transformers due to its simplicity and self-supervised nature. However, MLM operates at the token level and does not explicitly incorporate molecular properties or structural information into the learned representations.

To address this limitation, ChemBERTa-2 [1] has investigated chemically informed objectives via Multi-Task Regression (MTR) over molecular descriptors. This pretraining objective has shown to inject chemical knowledge into the models, improving downstream performance compared to MLM in downstream tasks. Other approaches use structural information, such as graph-based pretraining [22] or fingerprint supervision [27], yet these techniques are underexplored in the context of encoder-based molecular foundational models for SMILES.

Contrastive learning has also gained attention in molecular representation learning, typically relying on data augmentations or binary positive and negative pairs [27]. However, such formulations often discard the continuous similarity information available from molecular descriptors and fingerprints.

### 2.3 Graph-based and 3D-aware Molecular Representations

Alongside SMILES-based approaches, a parallel line of research has explored richer molecular modalities for representation learning. Graph neural network-based methods represent molecules as 2D graphs, where atoms are nodes and bonds are edges, enabling message-passing mechanisms that capture local chemical neighborhoods [22, 29]. More recently, 3D-aware models have incorporated geometric information such as atomic coordinates, achieving strong results on tasks where conformational properties are critical [10, 32].

Despite their effectiveness, graph-based and 3D-aware methods depend on the availability of molecular graph representations or pre-computed 3D conformers. These are not standard in large-scale databases, where compounds are typically stored as SMILES strings, making their application in high-throughput pipelines impractical.

## 3 MolDeBERTa Framework

In this section, we present the MolDeBERTa architectures, tokenization process, pretraining objectives, and pretraining data. The pipeline of the MolDeBERTa framework is illustrated in Figure 1.

**Figure 1:**
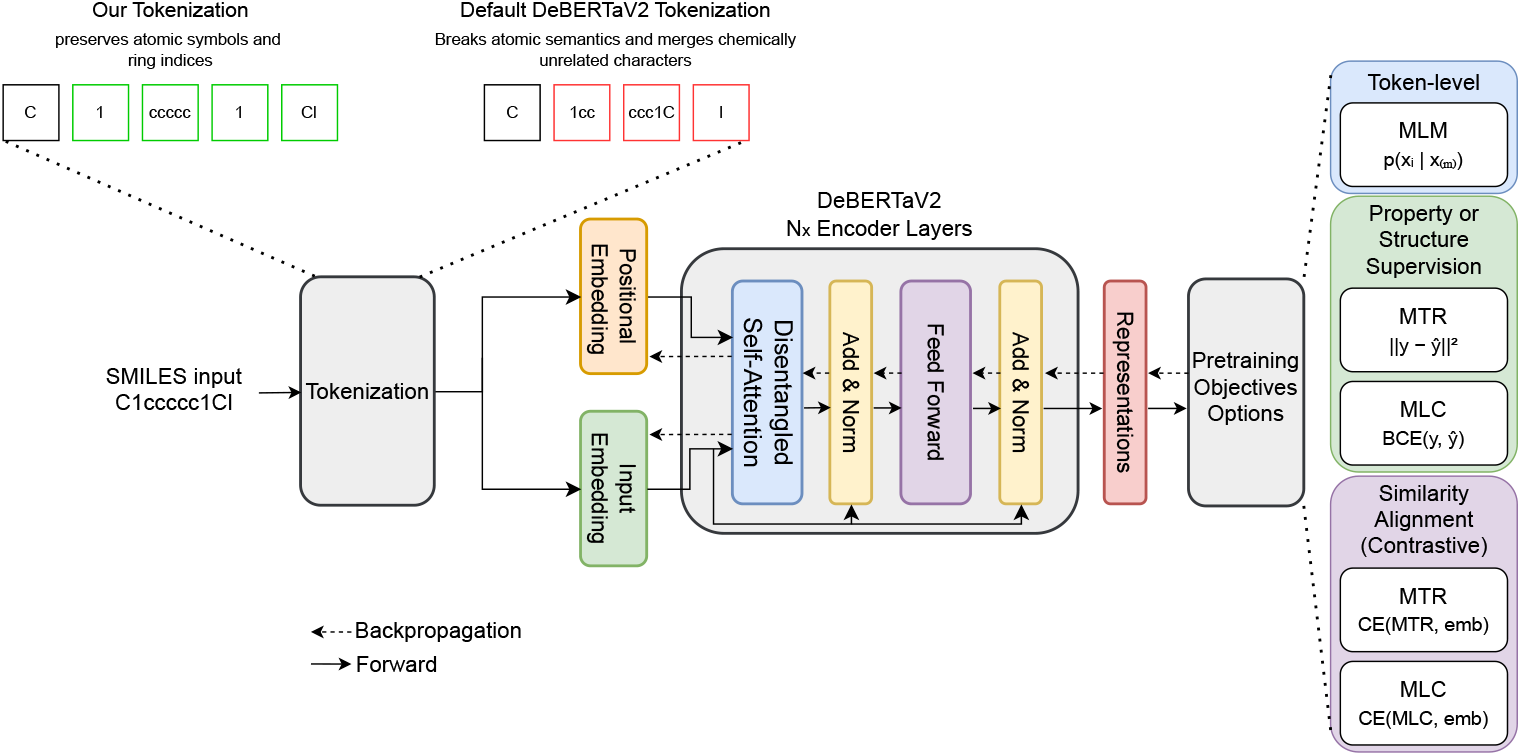
MolDeBERTa framework pipeline. Molecules in SMILES format are first tokenized using byte-level BPE and encoded by a DeBERTaV2 encoder. Byte-level BPE tokenization preserves atomic symbols and ring indices that encode chemical structure, whereas SentencePiece, used in standard DeBERTaV2, may merge chemically unrelated characters into subword units, disrupting SMILES syntax (top example of these two tokenization processes). We pretrained different configurations of the DeBERTaV2 architecture. During pre-training, we evaluated five training objectives, including token-level and chemistry-informed tasks, propagating the calculated error via backpropagation. We also investigated the impact of dataset size on pre-training performance. BCE and CE denote binary cross-entropy and cross-entropy losses, respectively, and emb denotes the learned molecular embedding.

### 3.1 MolDeBERTa Architecture

The MolDeBERTa framework is built upon the encoder-based DeBERTaV2 [9] transformer architecture, which has demonstrated strong performance across a wide range of NLP tasks. To evaluate the impact of model capacity, we instantiate MolDeBERTa at three different scales, i.e., tiny, small, and base. These variants differ in the number of encoder layers, hidden dimensions, and attention heads, while preserving the same overall architectural design. Table 1 summarizes the configuration of each model.

**Table 1:** Number of encoder layers, hidden dimension, and attention heads per encoder layer for each MolDeBERTa configuration.

| Architecture | Layers | Hidden Dimension | Attention Heads |
| --- | --- | --- | --- |
| Base | 12 | 768 | 12 |
| Small | 6 | 512 | 8 |
| Tiny | 3 | 384 | 6 |

Across all variants, MolDeBERTa follows a standard encoder-based transformer design. The maximum positional embedding length is set to 128 tokens, which covers the vast majority of molecules in our datasets while enabling efficient large-scale training. All architectural variants share the same tokenization scheme, positional encoding strategy, and training pipeline, enabling fair comparisons across model scales and pretraining objectives.

### 3.2 MolDeBERTa Tokenization

For tokenization, we employ a byte-level Byte-Pair Encoding (BPE) [24] tokenizer to process SMILES strings. In byte-level BPE, each input sequence is first decomposed into a sequence of raw bytes, i.e., ASCII characters, and subword units are then learned by iteratively merging frequent byte-level patterns. This design ensures that every character in the input is representable and prevents the generation of out-of-vocabulary tokens.

Unlike the original DeBERTaV2 architecture, which relies on SentencePiece [13] tokenization, we adopt byte-level BPE to better accommodate the character-level syntax of SMILES representations. SMILES strings contain a wide range of special characters (e.g., brackets, digits, bond symbols) whose semantics are sensitive to exact character composition. SentencePiece tokenization may fragment or merge character sequences in ways that obscure these syntactic constraints, whereas byte-level BPE preserves fine-grained character information while still learning frequent chemically meaningful substructures.

The tokenizer was trained on the entire 10M pretraining corpus (described in Section 3.4), using a vocabulary size of 4,000 tokens and a minimum token frequency of 2. This configuration balances expressiveness and vocabulary compactness while avoiding excessive fragmentation of chemically meaningful substructures. During tokenization, each SMILES sequence is prepended with a special classification token ([CLS]) and terminated with a separator token ([SEP]). The [CLS] token serves as a global aggregation point for the entire molecular sequence and is used for pretraining objectives and downstream tasks, while the [SEP] token marks sequence boundaries and stabilizes training across variable-length inputs.

### 3.3 MolDeBERTa Pretraining Objectives

We evaluate five distinct pretraining objectives for MolDeBERTa, each designed to inject different inductive biases into the learned molecular representations. The first pretraining objective is Masked Language Modeling (MLM), the standard task for transformer-based language models and used as a baseline. Given an input SMILES sequence, we randomly mask 15% of the tokens and train the model to predict the original masked tokens based on the contextualized representations produced by the final layer. The training objective minimizes the negative log-likelihood of the masked tokens, as defined in Equation 1, where x denotes a tokenized SMILES sequence and ℳ is the set of masked token positions.

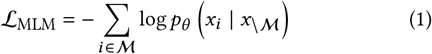

The second pretraining objective is Multi-Task Regression (MTR), which directly supervises the model using continuous molecular descriptors. Given a SMILES string, the model is trained to predict multiple real-valued molecular properties simultaneously. We employ the representation of the [CLS] token as a global molecular embedding and minimize the mean squared error between the predicted ýand target y descriptor values considering the n examples, as shown in Equation 2.

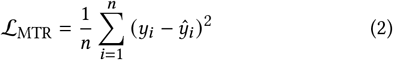

The third pretraining objective is Multi-Label Classification (MLC), which targets molecular substructure information. Each molecule is associated with a binary vector indicating the presence or absence of predefined substructures. Using the [CLS] token representation, the model is trained to predict this multi-label target via a binary cross-entropy loss, as defined in Equation 3, where n is the number of samples and m is the number of labels.

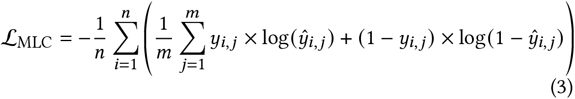

Finally, we consider contrastive variants of both MTR and MLC, referred as contrastive MTR and contrastive MLC. In these settings, we introduce a descriptor-informed contrastive objective that aligns the similarity structure induced by molecular descriptors with the similarity structure learned in the embedding space. Specifically, given a batch of *B* molecules, let *z*_*i*_ denote the embedding of molecule *i* obtained from the [CLS] token, and let y_*i*_ denote its corresponding molecular descriptor vector. We first compute pairwise cosine similarities in both spaces following Equation 4. Next, the descriptor-based similarities define a target distribution over molecular relationships via a temperature-scaled softmax, while the model induces a predicted distribution from the embedding similarities, following Equation 5. Finally, the contrastive objective minimizes the cross-entropy between these two distributions (Equation 6). During pretraining, the temperature parameter *τ* is learned jointly with the model parameters and is initialized to 0.1.

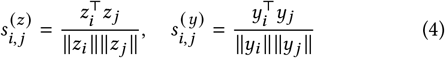

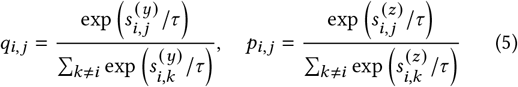

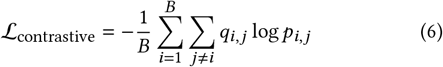

For pretraining objectives based on molecular properties and substructure, including both standard and contrastive variants, we derive all features directly from SMILES strings using the RDKit library [14]. Specifically, for MTR-based pretraining, we employ 216 real-valued molecular descriptors capturing physicochemical properties of molecules. For MLC-based pretraining, we utilize 2048-dimensional Morgan fingerprints, where each binary value indicates the presence or absence of a specific molecular substructure, generated with a radius of 2 around each atom. These descriptors and fingerprints are fully deterministic and reproducible from SMILES strings, requiring no additional experimental measurements or external annotations.

### 3.4 Pretraining Data

To pretrain the MolDeBERTa models, we utilize two datasets with substantially different scales. The first dataset, referred as 10M, corresponds to the dataset originally used for ChemBERTa-1 [6] pretraining and consists of 10 million molecules sampled from Pub-Chem [11]. The second dataset, referred to as 123M, comprises the complete PubChem collection, with 123 million molecules, obtained in December 2025. The dataset was used as provided by PubChem, without additional chemical normalization or canonicalization, and organized into a large-scale SMILES corpus suitable for transformer-based pretraining. To ensure reproducibility, the exact dataset used in this work is publicly released via Hugging Face^1^.

The pretraining is conducted using 99% of the data for training and 1% for validation. Since MTR-based objectives rely on continuous-valued targets, all molecular descriptors are normalized using the mean and standard deviation computed from the training set. Models pretrained with the MLM, MTR, and MLC objectives are optimized for 5 epochs. For contrastive MLC and contrastive MTR pretraining, we pretrain the models for 10 epochs on the 123M dataset and for 20 epochs on the 10M dataset. The number of training epochs was selected based on empirical convergence behavior observed during training, where contrastive objectives were found to converge significantly faster at larger dataset scales.

Details regarding the learning rate and batch size for each architecture are reported in Table 2. Batch sizes are chosen to maximize computational utilization while respecting memory constraints for each model size. All pretraining experiments were conducted using mixed-precision (float16) training on a cluster of six NVIDIA A6000 GPUs with AdamW [19] optimizer.

**Table 2:** Batch size and learning rate used for each model architecture during pretraining.

| Architecture Size | Batch Size | Learning Rate |
| --- | --- | --- |
| Base | 512 | 5e-5 |
| Small | 1024 | 5e-5 |
| Tiny | 2048 | 1e-4 |

## 4 Experimental Setup

### 4.1 Downstream Tasks

To evaluate MolDeBERTa framework and compare it with state-of-the-art, we benchmark our framework on nine datasets from the MoleculeNet [28] datasets. These datasets are widely adopted in the molecular machine learning literature and cover a diverse range of chemical properties, biological activities, and dataset scales, ranging from approximately 1,000 to 40,000 molecules. Specifically, we consider aqueous solubility (Delaney), lipophilicity (Lipo), inhibitory potency against the BACE-1 enzyme (BACE classification and BACE regression), hepatic drug clearance in humans (Clearance), blood-brain barrier permeability (BBBP), compound toxicity based on clinical trial data (ClinTox), inhibition of HIV replication (HIV), and activation of the p53 stress response pathway (Tox21). Among these, four datasets correspond to regression tasks (BACE, Clearance, Delaney, and Lipo), while five correspond to binary classification tasks (BACE, BBBP, ClinTox, HIV, and Tox21).

### 4.2 Comparison with State-of-the-Art Methods

To benchmark MolDeBERTa against the state-of-the-art in molecular representation learning, we evaluate our models against three categories of baselines. For SMILES-based encoders, we compare against ChemBERTa-1 [6]^2^, seven variants of ChemBERTa-2 [1], and MolFormer-XL [23]^3^. ChemBERTa-2 variants include ChemBERTa-2-5M-MLM^4^, ChemBERTa-2-5M-MTR^5^, ChemBERTa-2-10M-MLM^6^, ChemBERTa-2-10M-MTR^7^, ChemBERTa-2-77M-MLM^8^, ChemBERTa-2-77M-MTR^9^, and ChemBERTa-2-100M-MLM^10^.

For classical fingerprint-based baselines, we evaluate Random Forest models trained on RDKit molecular descriptors and Morgan fingerprints. For graph and 3D-aware models, we evaluate D-MPNN [29], a message-passing neural network trained from scratch on each downstream task, Uni-Mol [32] and Uni-Mol2 [10], 3D molecular encoders pretrained on molecular conformations. For Uni-Mol2, we use the 84M parameter variant, which is comparable in scale to the SMILES-based encoders evaluated in this work.

All SMILES-based models are initialized from their publicly released pretrained checkpoints on HuggingFace and fine-tuned using the same protocol applied to MolDeBERTa, ensuring a fair and controlled comparison. Methods for which pretrained weights are not publicly available, such as SMILES-BERT, are excluded from the evaluation due to reproducibility constraints. For MolFormer-XL, we evaluate the publicly available checkpoint pretrained on 10% of the data described in the original paper, as the full-data version is not publicly released. For Uni-Mol^11^, Uni-Mol2 and D-MPNN^12^, we apply their respective native procedures.

### 4.3 Fine-tuning Protocol

To ensure a fair and rigorous evaluation, we fine-tuned all 30 MolDeBERTa variants, and all baseline models, including SMILES-based encoders, Random Forest, D-MPNN, Uni-Mol, and Uni-Mol2, using a unified protocol, following ChemBERTa-2 [1] study. Hyperparameter optimization is performed with Optuna [2] over 20 trials, searching over learning rate, batch size, and random seed. Each trial is trained for up to 100 epochs with early stopping on validation loss, and the best validation checkpoint is evaluated on the held-out test set.

For all datasets, we utilize the scaffold split protocol implemented in DeepChem [21], which partitions molecules according to their scaffolds to prevent structural information leakage between training and evaluation sets. Each dataset is split into 80% training, 10% validation, and 10% test sets. For classification tasks, class imbalance is addressed by applying class-weighted loss functions. For regression tasks, target values are normalized to zero mean and unit variance during training, rescaling the predictions to the original space for evaluation. The performance is assessed using the area under the ROC curve (ROC-AUC) for classification tasks and the root mean squared error (RMSE) for regression tasks.

## 5 Experimental Results

### 5.1 Main Benchmark Results

In the primary evaluation of MolDeBERTa, we compare the best-performing configuration across all pretraining objectives, model architectures, and pretraining dataset scales against state-of-the-art models on nine downstream tasks from MoleculeNet. Table 3 reports the test-set performance, where the best result for each task is shown in bold and the second-best result is underlined.

**Table 3:** Test-set performance on four regression and five classification tasks from MoleculeNet. MolDeBERTa results correspond to the best-performing configuration selected among all pretraining objectives, model architectures, and pretraining dataset scales. Lower RMSE indicates better performance for regression tasks, while higher ROC-AUC indicates better performance for classification tasks. Best results are shown in bold and second-best results are underlined.

| Model | Regression (RMSE) |  |  |  | Classification (ROC-AUC) |  |  |  |  |
| --- | --- | --- | --- | --- | --- | --- | --- | --- | --- |
|  | BACE | Clearance | Delaney | Lipo | BACE | BBBP | ClinTox | HIV | Tox21 |
| MolDeBERTa (Our model) | **0.654** | **0.797** | 0.400 | <u>0.545</u> | 0.825 | **0.783** | **0.999** | 0.781 | <u>0.774</u> |
| ChemBERTa-1 | 0.760 | 1.085 | 0.495 | 0.695 | 0.774 | 0.753 | <u>0.997</u> | 0.767 | 0.669 |
| ChemBERTa-2-5M-MLM | 0.899 | 1.151 | 0.476 | 0.671 | 0.794 | 0.742 | 0.994 | 0.732 | 0.706 |
| ChemBERTa-2-5M-MTR | 0.745 | 1.133 | <u>0.394</u> | 0.572 | 0.821 | 0.735 | 0.992 | 0.756 | 0.734 |
| ChemBERTa-2-10M-MLM | 0.793 | 1.104 | 0.432 | 0.642 | 0.825 | 0.746 | 0.994 | 0.758 | 0.690 |
| ChemBERTa-2-10M-MTR | 1.012 | 1.092 | 0.396 | 0.574 | 0.804 | 0.724 | 0.993 | 0.751 | 0.753 |
| ChemBERTa-2-77M-MLM | 0.928 | 1.054 | 0.538 | 0.633 | <u>0.842</u> | 0.752 | 0.991 | 0.726 | 0.687 |
| ChemBERTa-2-77M-MTR | 1.077 | 1.091 | **0.384** | 0.577 | 0.783 | 0.695 | 0.988 | 0.733 | 0.693 |
| ChemBERTa-2-100M-MLM | 0.739 | 1.027 | 0.453 | 0.624 | 0.812 | 0.747 | 0.996 | 0.728 | 0.605 |
| MolFormer-XL | 0.785 | <u>0.946</u> | 0.431 | 0.547 | 0.824 | 0.740 | 0.996 | 0.694 | 0.593 |
| RF + RDKit | 0.803 | 1.011 | 0.458 | 0.631 | 0.818 | <u>0.761</u> | 0.856 | 0.774 | **0.777** |
| RF + FP | 0.928 | 1.063 | 0.798 | 0.718 | **0.860** | 0.686 | 0.812 | **0.805** | 0.748 |
| Uni-Mol | 0.830 | 0.987 | 0.426 | 0.554 | 0.754 | 0.679 | 0.735 | 0.725 | 0.711 |
| Uni-Mol 2 (84 M) | <u>0.730</u> | 0.992 | 0.423 | **0.509** | 0.815 | 0.716 | 0.910 | 0.705 | 0.750 |
| D-MPNN | 0.771 | 1.046 | 0.517 | 0.570 | 0.815 | 0.730 | 0.875 | <u>0.797</u> | 0.731 |

Overall, MolDeBERTa achieves the best overall performance on 4 out of 9 downstream tasks and outperforms all SMILES-based encoders on 7 out of 9 tasks. For regression tasks, MolDeBERTa exhibits strong improvements on BACE and Clearance, reducing the RMSE by approximately 10% compared to Uni-Mol2 and 16% compared to MolFormer-XL, respectively. In classification tasks, MolDeBERTa achieves consistent results on BBBP and ClinTox datasets, improving ROC-AUC by 2.2 percentage points on BBBP scenario.

Compared to classical models, MolDeBERTa outperforms both RF versions across 6 out of 9 tasks. Notably, RF with fingerprints achieves strong results in HIV and BBBP classification tasks, consistent with prior findings showing that fingerprint-based models can be effective for specific property prediction tasks [29]. Against graph-based models, MolDeBERTa consistently surpasses D-MPNN across 8 out of 9 benchmarks despite D-MPNN operating directly on 2D molecular graphs. This suggests that large-scale pretraining compensates for the lack of explicit graph structure in our approach. Compared to Uni-Mol and Uni-Mol2, which leverage explicit 3D conformational information, MolDeBERTa achieves superior performance on 8 out of 9 tasks, with particularly large margins on ClinTox and BBBP. These results demonstrate that chemistry-informed SMILES-based pretraining can match or surpass models that rely on explicit 3D geometry, while remaining significantly more scalable and accessible.

### 5.2 Effect of Pretraining Objective

We next analyze the impact of different pretraining objectives on downstream performance. To isolate the effect of the pretraining objective itself, we compare models with identical architectures and pretraining dataset sizes while varying only the objective. The complete quantitative results for all pretraining objectives and model architectures are reported in appendix. Here, we focus on the dominant trends and their implications.

Across all model scales and dataset sizes, we observe a similar pattern, i.e., pretraining objectives that directly supervise molecular properties or substructure information outperform MLM on the majority of downstream tasks. While MLM provides a generic learning by reconstructing masked SMILES tokens, it primarily captures syntactic regularities and does not explicitly encode physicochemical or functional molecular information. In contrast, MTR and MLC introduce chemical-related learning by directly supervising molecular descriptors and substructure fingerprints, respectively. As a result, models pretrained with MTR or MLC consistently outperform their MLM counterparts across most benchmarks.

We further observe an alignment between the pretraining and the downstream task type considering MTR and MLC objectives. MTR is particularly effective for regression benchmarks, where the targets correspond to continuous physicochemical properties, similar to the supervision features used during pretraining. For example, under the base architecture, MTR achieves the lowest RMSE in three out of four regression tasks compared to MLM and MLC, reducing the error by up to approximately 10% on the Lipo and Clearance datasets. Conversely, MLC pretraining tends to perform better on classification tasks, which are closely related to the presence or absence of molecular substructures encoded by fingerprint-based labels. In the same setting (base architecture), MLC improves ROC-AUC on the Tox21 benchmark by approximately 6.5% compared to both MTR and MLM. Interestingly, this direct alignment becomes less pronounced for the contrastive variants of MTR and MLC. By using molecular descriptors only to define similarity relationships rather than as explicit prediction targets during the pretraining step, contrastive pretraining emphasizes the organization of the embedding space rather than direct label reconstruction. As a result, for small architecture, contrastive MTR and contrastive MLC achieve the best performance across a broader set of tasks, including Clearance, Lipo, BBBP, ClinTox, and HIV, surpassing MLM as well as their non-contrastive counterparts. With that, this results in representations that are less related to a specific task type, but more robust and transferable across heterogeneous downstream objectives.

Our results demonstrate that the choice of pretraining objective plays a central role in molecular representation learning. Objectives that incorporate chemically meaningful supervision, either through direct prediction or descriptor-informed contrastive alignment, lead to molecular representations that generalize more effectively than those learned through generic token-level language modeling alone.

### 5.3 Effect of Model Scale

Then, we investigate the effect of model scale on downstream performance by comparing MolDeBERTa variants, i.e., tiny, small, and base. The complete quantitative results for all architectures are reported in the appendix.

Across all pretraining objectives and dataset scales, we observe a consistent improvement in downstream performance as model capacity increases. In general, base models outperform small models, which in turn outperform tiny models, with scaling leading to relative RMSE reductions up 20% on regression tasks and absolute gains up to 2 ROC-AUC points on classification benchmarks, indicating that larger architectures learn more expressive and transferable molecular representations.

However, the benefits of scaling are not uniform across pretraining objectives. For both tiny and base architectures, the best down-stream performance is typically achieved by models pretrained with MTR-based objectives, including both the standard and contrastive variants. Specifically, for the base architecture, models pretrained with MTR and contrastive MTR achieve the best results on 7 out of 9 tasks. Similarly, for the tiny architecture, MTR-based objectives account for 7 out of 9 best-performing configurations. In contrast, the small architecture exhibits a more dispersed performance profile across pretraining objectives, without a single objective consistently dominating across tasks. In this setting, no single objective consistently dominates, MLC and contrastive MLC account for 5 best results, while MTR and contrastive MTR account for 4 tasks.

Based on the results, we conclude that model size plays an important role in molecular representation learning. Increasing the model capacity consistently improves performance, particularly when combined with chemical-based pretraining objectives, highlighting the importance of jointly considering architecture scale and pretraining strategy.

### 5.4 Effect of Dataset Scale

Finally, we evaluate the impact of pretraining dataset scale on down-stream performance by comparing MolDeBERTa models pretrained on the 10M and 123M dataset. To isolate the effect of dataset size, we compare models with identical architectures and pretraining objectives while varying only the number of molecules used during pretraining. Detailed results are reported in the appendix.

The outcomes show that pretraining on the larger dataset present improvements across most downstream tasks and model configurations. In regression benchmarks, pretraining on 123M molecules yields RMSE reductions of approximately up to 10% compared to models pretrained on 10M molecules, with particularly consistent improvements observed for tasks such as BACE and Clearance. For classification benchmarks, larger-scale pretraining typically results in absolute ROC-AUC gains of up to 5 points, such as HIV task considering small architecture with contrastive MLC pretraining objective. This trend indicates that increased data diversity enables the model to learn richer and more generalizable molecular representations. However, we also observe that the benefits of increased dataset scale are not uniform across all settings. In some cases, models pretrained on the 10M dataset achieve superior performance on specific downstream tasks, such as Tox21 classification task using small architecture with MLC pretraining objective. This behavior suggests that in such cases the model has already captured the general context of molecules and additional data can introduce redundancy rather than novel learning information.

### 5.5 Chemistry-Informed Objectives Generalize Across Architectures

To assess whether the benefits of chemistry-informed pretraining are specific to the DeBERTaV2 backbone or generalize across encoder architectures, we apply all five pretraining objectives to the ChemBERTa-2 architecture, a RoBERTa-based encoder, using the 10M dataset. This provides a controlled experiment where the architecture is replaced while the pretraining protocol remains identical. Although public checkpoints for ChemBERTa-2 (MLM and MTR) exist, including a version pretrained on 10 million SMILES, the unavailability of the exact data and original training code precludes a strictly controlled comparison. Therefore, to ensure a fair evaluation where the pretraining objective is the sole variable, we pretrained the ChemBERTa-2 architecture from scratch, initializing with random weights, across all five objectives using our protocol.

Full downstream performance metrics for all configurations are detailed in Table 4. Our results show that chemistry-informed objectives yielded consistent gains over the standard MLM baseline. Most notably, on the Tox21 benchmark, contrastive-based MTR achieved an ROC-AUC of 0.763, outperforming the standard MLM baseline, which achieved a ROC-AUC value of 0.678. Furthermore, for regression tasks such as Lipo, the explicitly structural guidance of contrastive-based MLC proved superior, lowering the RMSE from 0.647 of MLM-based pretraining to 0.622.

**Table 4:** Test-set performance on four regression and five classification tasks from ChemBERTa-2 pretrained with different objectives on 10M dataset. Lower RMSE indicates better performance for regression tasks, while higher ROC-AUC indicates better performance for classification tasks.

| Pretraining Objective | Regression (RMSE) |  |  |  | Classification (ROC-AUC) |  |  |  |  |
| --- | --- | --- | --- | --- | --- | --- | --- | --- | --- |
|  | BACE | Clearance | Delaney | Lipo | BACE | BBBP | ClinTox | HIV | Tox21 |
| MLM | 0.812 | 1.121 | 0.458 | 0.647 | 0.805 | 0.724 | 0.993 | 0.725 | 0.678 |
| MLC | 0.853 | 1.180 | 0.528 | 0.764 | 0.760 | 0.728 | 0.993 | 0.776 | 0.662 |
| Contrastive MLC | 0.777 | 1.096 | 0.454 | 0.622 | 0.795 | 0.767 | 0.991 | 0.733 | 0.732 |
| MTR | 0.892 | 1.100 | 0.443 | 0.633 | 0.748 | 0.735 | 0.996 | 0.735 | 0.656 |
| Contrastive MTR | 0.886 | 1.116 | 0.417 | 0.632 | 0.812 | 0.742 | 0.993 | 0.768 | 0.763 |

These results demonstrate that chemistry-informed pretraining objectives (MTR, MLC, and their contrastive variants) yield consistent improvements over MLM regardless of the underlying encoder architecture. This suggests that the benefits of chemistry-informed supervision are architecture-agnostic and transferable to other SMILES-based encoders beyond MolDeBERTa.

## 6 Interpretability and Scientific Validation

Beyond the predictive performance, an important aspect for molecular foundational models is that their learned representations align with established chemical principles. To assess whether MolDeBERTa captures chemically meaningful patterns, we conduct a qualitative interpretability analysis on downstream regression tasks using gradient-based attribution methods. As a case study, we analyze attribution patterns for ibuprofen for Delaney (aqueous solubility) and Lipo (lipophilicity) datasets. This molecule is particularly interesting for this analysis, as its molecular structure contains both a hydrophobic aromatic backbone and a polar carboxylic acid group, each contributing differently to aqueous solubility and lipophilicity.

All interpretability experiments were performed using the MolDeBERTa base architecture pretrained with the MTR objective on the 123M dataset, which consistently achieved strong performance across downstream benchmarks. The pretrained model was fine-tuned separately for each downstream task. To quantify token importance, we employ SHAP technique [20], which computes shapley value attributions by integrating gradients along a path between a baseline input and the actual input. The resulting attribution scores reflect the contribution of each input token to the model’s prediction. To enable molecular-level interpretation, we map token-level attributions back to individual atoms using an alignment between SMILES tokens and atomic symbols.

Figure 2 illustrates the attribution maps for ibuprofen. Warmer colors indicate higher attribution values, while cooler colors indicate lower importance. For aqueous solubility prediction, the model assigns the highest attribution to the carboxylic acid group, which is consistent with its ability to form hydrogen bonds and increase solubility in water. On the other hand, for lipophilicity prediction, attribution shifts toward the carbon backbone, reflecting the dominant contribution of hydrophobic regions to lipid solubility. This contrast demonstrates that, despite sharing the same pretrained backbone, MolDeBERTa learns task-specific molecular representations during fine-tuning on different downstream tasks that align with known physicochemical mechanisms.

**Figure 2:**
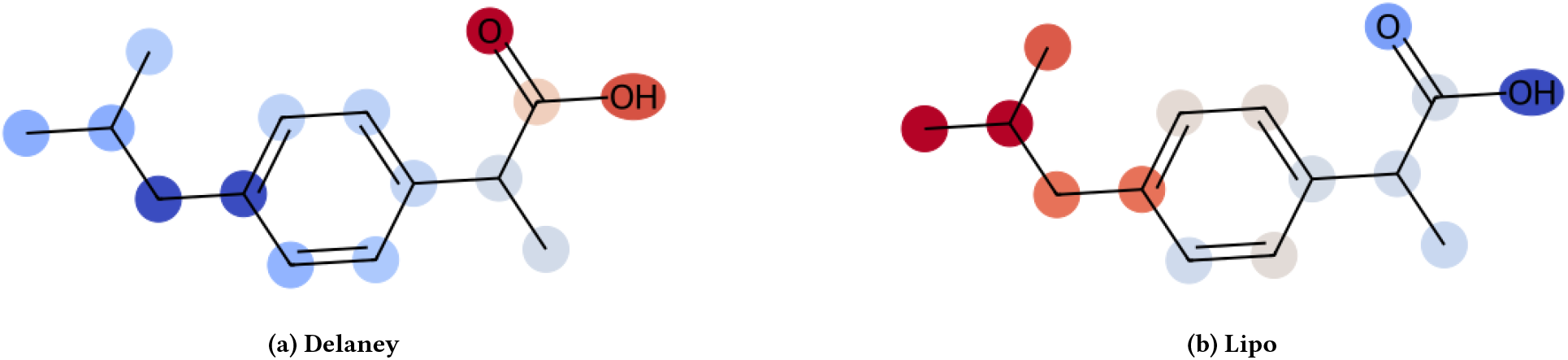
Atom-level attribution maps for ibuprofen considering SHAP attributions for MolDeBERTa on Delaney and Lipo datasets. Red denotes higher importance and blue lower importance. The model focuses on hydrophobic regions for lipophilicity and on the polar carboxylic acid group for aqueous solubility, reflecting task-specific chemical relevance.

## 7 Limitations

Despite the strong empirical performance demonstrated by MolDeBERTa, several limitations should be acknowledged. First, the model operates exclusively on SMILES-based textual representations and does not explicitly incorporate three-dimensional molecular geometry. As a result, MolDeBERTa can exhibit degraded performance on tasks that critically depend on 3D conformational information or stereochemistry. Second, the models are pretrained on small molecules from PubChem and rely on a maximum input length of 128 tokens. Consequently, representations of large or highly complex molecules may be truncated or inadequately captured, potentially limiting performance for such compounds. Third, the descriptor-based pretraining objectives rely on molecular properties and fingerprints computed using RDKit. While these descriptors are widely used and efficiently computed, they represent heuristic approximations of molecular behavior. As a result, the learned representations may be biased toward the assumptions and coverage of these descriptors and may not fully capture rare or underrepresented molecular properties. Fourth, our experimental evaluation is restricted to datasets from the MoleculeNet benchmark. Although these datasets span a broad range of molecular prediction tasks and are widely adopted in the literature, performance can vary on other benchmarks or real-world industrial datasets. Extending evaluation to additional domains is an important direction for future work.

## 8 Conclusions

We designed, developed, trained and evaluated *MolDeBERTa*, an encoder-based molecular foundational model. *MolDeBERTa* uses DeBERTaV2 backbone architecture and pretrained on up to 123M SMILES from PubChem with chemistry-informed objectives, including multi-label substructure classification and descriptor-informed contrastive learning. These objectives explicitly align the latent space with physicochemical properties *without* requiring experimental labels or 3D geometry.

MolDeBERTa achieves state-of-the-art performance on 4 out of 9 MoleculeNet benchmarks, outperforming SMILES-based encoders, fingerprint-based models, graph neural networks, and 3D molecular models under a unified fine-tuning protocol. Our experiments demonstrate that chemistry-informed pretraining objectives substantially outperform MLM, that contrastive variants further improve representation organization, and that scaling benefits depend on the interaction between architecture size and the objective. Atom-level interpretability analyses confirm that MolDeBERTa learns task-specific representations consistent with known physicochemical principles.

Future work includes extending MolDeBERTa to additional molecular tasks and integrating 3D structural information to further enhance representation learning. All pretrained models, datasets, and code are publicly available to promote reproducibility.

## Acknowledgments

Research reported in this publication was supported by NIGMS of the National Institutes of Health under award number R35GM153434. The content is solely the responsibility of the authors and does not necessarily represent the official views of the National Institutes of Health.

## A Detailed Benchmark Results

Tables 5, 6, and 7 report the complete downstream performance for tiny, small, and base architectures, respectively.

**Table 5:** Test-set performance on four regression and five classification tasks from MoleculeNet considering tiny architecture. Lower RMSE indicates better performance for regression tasks, while higher ROC-AUC indicates better performance for classification tasks.

| Pretraining Objective | Pretraining Dataset | Regression (RMSE) |  |  |  | Classification (ROC-AUC) |  |  |  |  |
| --- | --- | --- | --- | --- | --- | --- | --- | --- | --- | --- |
|  |  | BACE | Clearance | Delaney | Lipo | BACE | BBBP | ClinTox | HIV | Tox21 |
| MLM | 123M | 0.873 | 1.080 | 0.485 | 0.651 | 0.784 | 0.737 | 0.997 | 0.735 | 0.713 |
| MLM | 10M | 0.892 | 1.156 | 0.492 | 0.649 | 0.760 | 0.724 | 0.996 | 0.773 | 0.687 |
| MLC | 123M | 0.777 | 1.084 | 0.521 | 0.642 | 0.784 | 0.723 | 0.995 | 0.763 | 0.722 |
| MLC | 10M | 0.835 | 1.018 | 0.477 | 0.641 | 0.813 | 0.753 | 0.988 | 0.760 | 0.756 |
| Contrastive MLC | 123M | 0.729 | 1.113 | 0.480 | 0.632 | 0.795 | 0.729 | 0.995 | 0.718 | 0.764 |
| Contrastive MLC | 10M | 0.804 | 1.105 | 0.517 | 0.655 | 0.808 | 0.731 | 0.995 | 0.769 | 0.705 |
| MTR | 123M | 0.770 | 1.005 | 0.471 | 0.606 | 0.770 | 0.765 | 0.999 | 0.764 | 0.748 |
| MTR | 10M | 0.783 | 1.035 | 0.454 | 0.643 | 0.784 | 0.756 | 0.986 | 0.776 | 0.700 |
| Contrastive MTR | 123M | 0.784 | 1.029 | 0.421 | 0.590 | 0.790 | 0.747 | 0.999 | 0.727 | 0.757 |
| Contrastive MTR | 10M | 0.757 | 1.045 | 0.417 | 0.634 | 0.796 | 0.750 | 0.996 | 0.763 | 0.735 |

**Table 6:** Test-set performance on four regression and five classification tasks from MoleculeNet considering small architecture. Lower RMSE indicates better performance for regression tasks, while higher ROC-AUC indicates better performance for classification tasks.

| Pretraining Objective | Pretraining Dataset | Regression (RMSE) |  |  |  | Classification (ROC-AUC) |  |  |  |  |
| --- | --- | --- | --- | --- | --- | --- | --- | --- | --- | --- |
|  |  | BACE | Clearance | Delaney | Lipo | BACE | BBBP | ClinTox | HIV | Tox21 |
| MLM | 123M | 0.817 | 0.934 | 0.469 | 0.597 | 0.813 | 0.746 | 0.990 | 0.719 | 0.693 |
| MLM | 10M | 0.783 | 1.035 | 0.454 | 0.643 | 0.784 | 0.756 | 0.986 | 0.776 | 0.700 |
| MLC | 123M | 0.727 | 0.968 | 0.504 | 0.615 | 0.781 | 0.731 | 0.995 | 0.771 | 0.696 |
| MLC | 10M | 0.796 | 0.985 | 0.482 | 0.651 | 0.825 | 0.744 | 0.987 | 0.756 | 0.771 |
| Contrastive MLC | 123M | 0.750 | 1.018 | 0.452 | 0.570 | 0.825 | 0.743 | 0.998 | 0.781 | 0.737 |
| Contrastive MLC | 10M | 0.740 | 1.134 | 0.483 | 0.643 | 0.794 | 0.717 | 0.993 | 0.410 | 0.728 |
| MTR | 123M | 0.737 | 1.118 | 0.410 | 0.563 | 0.780 | 0.756 | 0.997 | 0.761 | 0.702 |
| MTR | 10M | 0.748 | 0.967 | 0.437 | 0.608 | 0.772 | 0.774 | 0.996 | 0.743 | 0.693 |
| Contrastive MTR | 123M | 0.784 | 0.889 | 0.434 | 0.545 | 0.786 | 0.776 | 0.996 | 0.732 | 0.734 |
| Contrastive MTR | 10M | 0.803 | 1.012 | 0.464 | 0.610 | 0.772 | 0.723 | 0.988 | 0.738 | 0.717 |

**Table 7:** Test-set performance on four regression and five classification tasks from MoleculeNet considering base architecture. Lower RMSE indicates better performance for regression tasks, while higher ROC-AUC indicates better performance for classification tasks.

| Pretraining Objective | Pretraining Dataset | Regression (RMSE) |  |  |  | Classification (ROC-AUC) |  |  |  |  |
| --- | --- | --- | --- | --- | --- | --- | --- | --- | --- | --- |
|  |  | BACE | Clearance | Delaney | Lipo | BACE | BBBP | ClinTox | HIV | Tox21 |
| MLM | 123M | 0.728 | 0.879 | 0.434 | 0.650 | 0.819 | 0.717 | 0.995 | 0.731 | 0.662 |
| MLM | 10M | 0.654 | 0.939 | 0.475 | 0.642 | 0.803 | 0.754 | 0.993 | 0.634 | 0.477 |
| MLC | 123M | 0.748 | 0.964 | 0.469 | 0.609 | 0.807 | 0.729 | 0.999 | 0.464 | 0.643 |
| MLC | 10M | 0.798 | 1.070 | 0.493 | 0.620 | 0.813 | 0.745 | 0.948 | 0.742 | 0.763 |
| Contrastive MLC | 123M | 0.763 | 0.933 | 0.470 | 0.600 | 0.795 | 0.751 | 0.991 | 0.751 | 0.774 |
| Contrastive MLC | 10M | 0.778 | 0.980 | 0.512 | 0.625 | 0.784 | 0.759 | 0.993 | 0.709 | 0.605 |
| MTR | 123M | 0.687 | 0.797 | 0.428 | 0.549 | 0.824 | 0.750 | 0.998 | 0.753 | 0.621 |
| MTR | 10M | 0.725 | 0.864 | 0.455 | 0.582 | 0.806 | 0.783 | 0.980 | 0.703 | 0.716 |
| Contrastive MTR | 123M | 0.658 | 1.121 | 0.419 | 0.566 | 0.802 | 0.757 | 0.980 | 0.753 | 0.659 |
| Contrastive MTR | 10M | 0.756 | 0.948 | 0.400 | 0.594 | 0.756 | 0.762 | 0.987 | 0.732 | 0.647 |

## Footnotes

1 https://huggingface.co/datasets/SaeedLab/MolDeBERTa

2 https://huggingface.co/seyonec/ChemBERTa-zinc-base-v1

3 https://huggingface.co/ibm-research/MoLFormer-XL-both-10pct

4 https://huggingface.co/DeepChem/ChemBERTa-5M-MLM

5 https://huggingface.co/DeepChem/ChemBERTa-5M-MTR

6 https://huggingface.co/DeepChem/ChemBERTa-10M-MLM

7 https://huggingface.co/DeepChem/ChemBERTa-10M-MTR

8 https://huggingface.co/DeepChem/ChemBERTa-77M-MLM

9 https://huggingface.co/DeepChem/ChemBERTa-77M-MTR

10 https://huggingface.co/DeepChem/ChemBERTa-100M-MLM

11 https://unimol.readthedocs.io

12 https://github.com/chemprop/chemprop

## Notes

### Competing Interest Statement

The authors have declared no competing interest.

### Summary of Updates

1. The title has been changed to correctly reflect the work. 2. Funding acknowledgement has changed - required by the federal agency. 3. Minor corrections to the text is done.

